# Immune Checkpoint Activity Regulates Polycystic Kidney Disease Progression

**DOI:** 10.1101/2022.04.24.489306

**Authors:** Emily K. Kleczko, Dustin T. Nguyen, Kenneth H. Marsh, Colin D. Bauer, Amy S. Li, Seth B. Furgeson, Berenice Y. Gitomer, Michel B. Chonchol, Eric T. Clambey, Kurt A. Zimmerman, Raphael A. Nemenoff, Katharina Hopp

**Author notes:** Corresponding Author: Dr. Katharina Hopp, Division of Renal Diseases & Hypertension University of Colorado Anschutz Medical Campus 12700 E 19th Avenue, P15-7400A, Mail Stop C281 Aurora, CO 80045. Contributed equally to this work (Emily K. Kleczko, Dustin T. Nguyen).

## Abstract

Innate and adaptive immune cells modulate Autosomal Dominant Polycystic Kidney Disease (ADPKD) severity, a common kidney disease with inadequate treatment options. ADPKD shares parallels with cancer where immune checkpoint inhibitors have been shown to reactivate CD8^+^ T cells and slow tumor growth. We have shown that, in PKD, CD8^+^ T cell loss worsens disease. This study used orthologous early-onset and adult-onset ADPKD models (*Pkd1* p.R3277C) to evaluate the role of immune checkpoints in PKD. Flow cytometry of kidney cells showed increased levels of PD-1 on CD8^+^ T cells and PD-L1 on macrophages and epithelial cells in *Pkd1*^RC/RC^ mice versus wildtypes, paralleling disease severity. PD-L1 was also upregulated in ADPKD human cells and patient kidney tissue versus controls. Genetic PD-L1 loss or treatment with an anti-PD-1 antibody did not impact PKD severity in early-onset or adult-onset ADPKD models. However, treatment with anti-PD-1 plus anti-CTLA-4, blocking two immune checkpoints, improved PKD outcomes in adult-onset ADPKD mice; neither monotherapy altered PKD. Combination therapy resulted in increased kidney CD8^+^ T cell numbers/activation and decreased kidney regulatory T cell numbers. Together, our data suggests that immune checkpoint activation is an important feature of and potential novel therapeutic target in ADPKD.

## INTRODUCTION

Autosomal Dominant Polycystic Kidney Disease (ADPKD), caused predominantly by mutations to *PKD1* or *PKD2,* is the most common monogenic kidney disease(1, 2). It is characterized by the growth of bilateral kidney cysts leading to end stage kidney disease (ESKD) in 50% of patients by middle age(3, 4). ADPKD patients present with substantial variability in disease severity and progression which cannot be solely explained by genic or allelic effects (5–7). Underlying reasons for this phenotypic heterogeneity remain poorly understood, but likely involve differences in the cystic microenvironment (CME). In cancer, a disease that shares multiple characteristics with PKD, innate and adaptive immune cells within the tumor microenvironment (TME) significantly modulate disease progression(8, 9). Similarly, studies of the CME suggest that immune cells play a key role in modulating PKD severity(10). Multiple groups have demonstrated that kidney macrophages drive cyst expansion in mouse models of ADPKD(11–18). Further, urinary CD4^+^ T cell numbers were reported to correlate with renal function loss in ADPKD patients, and we have shown that, correlative with disease severity, numbers of both CD4^+^ and CD8^+^ T cells increase in the cystic kidneys of an orthologous ADPKD mouse model and that depletion of CD8^+^ T cells resulted in more rapidly progressive cyst growth, suggesting an anti-cystogenic role of these cells(19, 20).

In cancer, tumor-specific naïve CD8^+^ T cells become activated by recognizing tumor specific neoantigens and self-antigens leading to production of cytotoxic molecules, such as granzymes and perforin, that directly kill tumor cells. Further, they secrete cytokines, such as IFNɣ, which increase expression of major histocompatibility complex (MHC) class I antigens by tumor cells, thereby rendering them better targets for CD8^+^ T cell-mediated killing(21, 22). A similar immunosurveillance role for CD8^+^ T cells can be proposed in PKD where CD8*^+^* T cell-recruitment and activation is a response to neoantigens produced by the de-differentiation process of the tubular epithelium during cyst development/growth (e.g., genomic instability, somatic mutations, damaged interstitial/epithelial cells). This correlates with our prior findings that CD8^+^ T cells localize to cystic lesions, that levels of IFNɣ are higher in cystic kidneys compared to controls, and that CD8^+^ T cell loss resulted in more rapidly progressive PKD(19).

Despite the presence of tumor-infiltrating effector CD8^+^ T cells most tumors continue to progress, suggesting that tumor reactive CD8^+^ T cells become dysfunctional, i.e. exhausted, during tumor progression(23, 24). This in large part, has been attributed to tumors being able to engage mechanisms that drive immunosuppression. Immunosuppressive cells that inhibit CD8^+^ T cell function include CD4^+^ T regulatory cells (T_Regs_), which inhibit CD8^+^ T cell activity via direct cytolysis or release of inhibitory cytokines (e.g., TGF-β, IL-10, IL-35), as well as M2-like tumor associated macrophages (TAMs) and myeloid-derived suppressor cells (MDSCs)(25). Both, MDSCs and TAMs, produce high levels of immunosuppressive proteins such as IL-10, TGF-β, IDO1, ARG1, iNOS and immune checkpoint ligands (e.g., PD-L1), causing CD8^+^ T cells to undergo metabolic reprogramming, lose effector function (secretion of TNF-α, IL-2, IFNɣ), and sustain expression of inhibitory receptors (PD-1, TIM-3, CTLA-4); all features of exhausted CD8^+^ T cells(21, 25). Interestingly, PKD-associated macrophages have been compared to TAMs due to multiple phenotypic and functional similarities(26).

Immune checkpoints, such as programmed cell death protein 1|programmed cell death ligand 1 (PD-1|PD-L1), are proteins that regulate immune homeostasis, prevent excessive CD8^+^ T cell activation, and protect against autoimmunity(27). Tumors have exploited this pathway to escape immune surveillance. In tumors, PD-1 is expressed on CD8^+^ T cells and binds to PD-L1 expressed on tumor cells and tumor infiltrating myeloid cells (e.g., TAMs). Interaction of these proteins dampens effector CD8^+^ T cell function and diminishes their ability to recognize and clear tumor cells(28). Immune checkpoint inhibitors (ICIs) are monoclonal antibodies that makeup a novel class of immunotherapy drugs that disrupt the interaction between ligand and receptor (e.g., PD-1 and PD-L1) leading to reactivation of anti-tumor T cells. PD-1|PD-L1 ICIs are FDA approved and have become the standard of care in multiple different cancer types, including renal cell carcinoma (RCC)(29–31).

A role for PD-1|PD-L1-associated immunomodulation has been suggested in various kidney pathologies. For example, PD-1 deficient mice develop lupus-like glomerulonephritis as they age (> 14 months of age) consistent with a role for immune checkpoints in self-tolerance(32). Consequently, adenovirus-driven PD-L1 overexpression improved lupus nephritis-associated pathologies in mice(33). A protective role of PD-1 has also been suggested in murine adriamycin nephropathy where treatment with anti-PD-1 led to worsened glomerular and tubulointerstitial injury(34). Contrastingly, a study found PD-1 expression to be significantly elevated on peripheral blood CD4^+^ T cells, CD8^+^ T cells, and CD19^+^ B cells of primary proliferative and non-proliferative glomerulonephritis patients versus controls and correlative with disease severity parameters, suggesting that ICIs may help glomerulonephritis patients(35). Similarly, increased PD-L1 expression on tubular epithelial cells was found in kidney biopsies of patients with IgA nephropathy, interstitial nephritis, and lupus nephritis compared to normal kidneys, although the functional relevance remains unclear(36).

Given the similarities between cancer and ADPKD, which has been described as a “neoplasia in disguise”, we examined whether targeting PD-1|PD-L1 represents a potential novel therapeutic target for treatment of ADPKD (8, 37). Using an orthologous model of ADPKD1, the *Pkd1* p.R3277C mouse(38), with varying rates of disease progression, we found a significant upregulation of both PD-1 on kidney CD8^+^ T cells as well as PD-L1 on kidney macrophages and epithelial cells. While genetic loss of PD-L1 or inhibition of PD-1 with a monoclonal antibody did not slow cyst progression in our models, combination treatment of α-PD-1 and a monoclonal antibody inhibiting a different immune checkpoint, cytotoxic T-lymphocyte-associated protein 4 (CTLA-4), significantly slowed PKD, suggesting, as seen in cancer clinical trials, that dual immune checkpoint blockade increases efficacy and response rate(39, 40). Our data also defines adaptive immunity pathways in the CME that are altered in response to these agents.

## RESULTS

### The PD-1|PD-L1 immune checkpoint pathway is induced in an orthologous mouse model of ADPKD

We analyzed the expression of PD-1 on kidney CD8^+^ T cells as well as PD-L1 on kidney epithelial cells and macrophages employing flow cytometry of kidney single cell suspensions obtained from an orthologous adult-onset ADPKD1 model, the *Pkd1*^RC/RC^ mouse(38). As previously published, the model presents with varying disease course in different mouse strains(19, 38, 41–43). C57Bl/6J *Pkd1*^RC/RC^ mice have gradually progressive PKD with the kidney area occupied by cysts increasing from ∼19% at 3 months of age to ∼23% at 9 months of age. Comparably, at 3 months of age the cystic kidney area of BALB/cJ *Pkd1*^RC/RC^ mice is ∼32% and of 129S6/SvEvTac *Pkd1*^RC/RC^ mice is ∼34%, highlighting a much more rapid PKD progression in these strains(19, 43).

We examined PD-1|PD-L1 expression in kidneys obtained from *Pkd1*^RC/RC^ mice and strain-matched wildtypes, male and female mice combined, at 3-, 6-, and 9-months of age to correlate the engagement of this immune checkpoint with severity of cystic kidney disease. Details on the PKD phenotype parameters of these mice have been previously published(19, 43). In all three strains, we found that expression of PD-1 on kidney CD8^+^ T cells in *Pkd1*^RC/RC^ mice increased correlative with cystic kidney disease severity compared to strain-, sex-, and age-matched wildtypes (**Figure 1A, B**). As previously published, C57Bl/6J wildtypes have more kidney CD8^+^ T cells compared to age-, sex-matched mice in the 129S6/SvEvTac or BALB/cJ strain(19). Correlatively, C57Bl/6J wildtypes had a high basal level of PD-1 positive CD8^+^ T cells, which, however, significantly increased in *Pkd1*^RC/RC^ mice at all three analyzed time points. This increase of PD-1 positive kidney CD8^+^ T cells paralleled the overall increase in CD8^+^ T cell numbers (**Supplemental Table 1**). Differently though, in the strains that present with more rapidly progressive PKD (129S6/SvEvTac or BALB/cJ), not only did PD-1 positive kidney CD8^+^ T cells increase in number with more severe cystic kidney disease when comparing *Pkd1*^RC/RC^ to wildtype mice, but PD-1^+^ CD8^+^ T cells became also more prominent among the CD8^+^ T cell population (**Figure 1A, B,Supplemental Table 1**).

**Figure 1.**
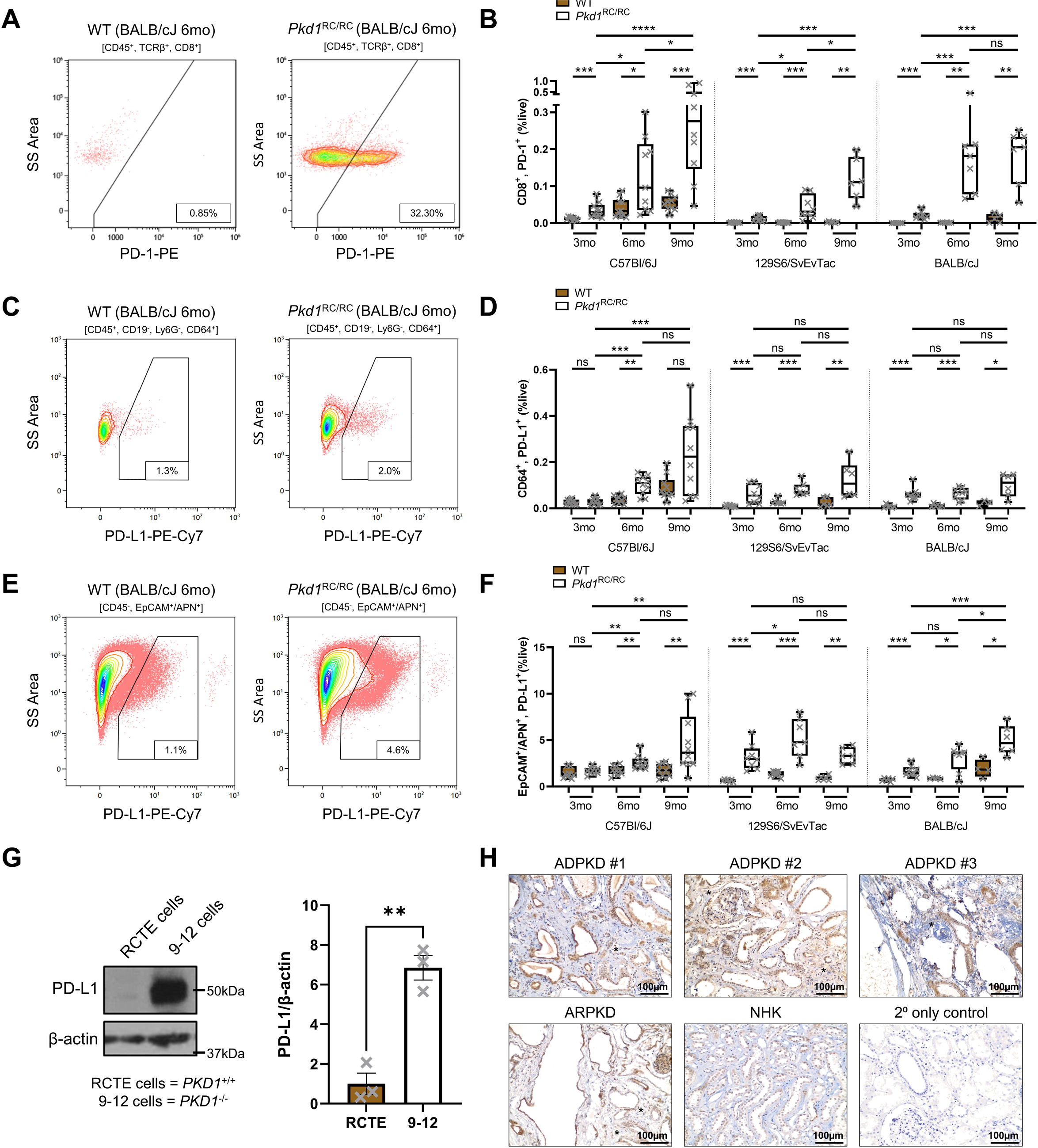
The PD-1/PD-L1 immune checkpoint pathway is induced in autosomal dominant polycystic kidney disease. Kidneys of wildtype (WT) and *Pkd1*^RC/RC^ mice in the C57Bl/6J, 129S6/SvEvTac, and BALB/cJ strains were harvested at 3, 6, and 9 months (mo) of age and analyzed by flow cytometry to identify the expression of PD-1 on CD8^+^ T cells (CD45^+^/TCRβ^+^/CD8^+^; **A, B**); PD-L1 on macrophages (CD45^+^/CD19^-^/Ly6G^-^/CD64^+^; **C, D**); and PD-L1 on epithelial cells (CD45^-^/EpCAM^+^/APN^+^; **D, F**). (**A, C, E**) Representative flow diagrams of 6-month-old BALB/cJ WT and *Pkd1*^RC/RC^ kidneys. Quantification of PD-1 positive CD8^+^ T cells (**B**), PD-L1 positive CD64^+^ macrophages (**D**), and PD-L1 positive epithelial cells (**F**) as percent live (%live) comparing WT (brown) to *Pkd1*^RC/RC^ (white) kidneys. PD-1 or PD-L1 expression is significantly upregulated on the respective cell type in *Pkd1*^RC/RC^ compared to WT kidneys and correlative with increasing disease severity in most instances. (**G**) PD-L1 expression in immortalized human renal cortical tubular epithelial (RCTE) cells (*PKD1*^+/+^) versus 9-12 cells (*PKD1*^-/-^). 9-12 cells have significantly increased levels of PD-L1 compared to control. Left panel: representative western blot image. Right panel: quantification of western blots from three independent samples. (**H**) Immunohistochemistry staining for PD-L1 in end-stage kidney tissue of three Autosomal Dominant Polycystic Kidney Disease (ADPKD) patients, an Autosomal Recessive PKD (ARPKD) patient, and a normal human kidney (NHK). 2° only control: kidney tissue slide of an ADPKD patient stained without adding primary antibody. The cystic epithelium in ADPKD and ARPKD shows increased expression of PD-L1 compared to PD-L1 expression in tubules of NHK. PD-L1 expression is also found in some interstitial cells within the ADPKD or ARPKD tissue sections (*). Data are presented as box plot with whiskers of 10-90th percentile (**B, D, F**) or the mean ± standard error of the mean (SEM, **G**); single data points are depicted. Non-parametric Mann-Whitney tests were performed. ns, non-significant. *p<0.05; **p<0.01; ***p<0.001; ****p<0.0001. N=9-10 for C57Bl/6J WT or *PKD1*^RC/RC^; N=6-8 for 129S6/SvEvTac WT or *PKD1*^RC/RC^; N=5-6 for BALB/cJ WT and N=7-8 BALB/cJ *PKD1*^RC/RC^. Data points are half male and half female for all groups.

It has been reported in multiple murine PKD models that the number of kidney macrophages increases in cystic kidneys compared to wildtype control kidneys(10). Correlatively, multiple studies have shown that depletion of kidney macrophages slows PKD progression in murine models(10). When evaluating kidney macrophage numbers (CD45^+^, CD64^+^ cells) in the *Pkd1*^RC/RC^ model compared to wildtype, we also found this population to increase in all three strains and correlatively with cystic kidney disease severity, although kidneys with the mildest cystic burden (C57Bl/6J *Pkd1*^RC/RC^ kidney at 3-and 6-months of age) showed only a moderate, non-significant increase in CD64^+^ kidney macrophage numbers compared to wildtype controls (**Supplemental Table 2**). Since TAMs have been found to be a prominent driver of immunosuppression and are known to express PD-L1 in cancer, we focused on the expression of PD-L1 on CD64^+^ kidney macrophages(44). We found a significant increase of PD-L1 positive kidney macrophages in *Pkd1*^RC/RC^ mice compared to wildtypes even in moderately cystic kidneys (e.g., 6-month-old C57Bl/6J *Pkd1*^RC/RC^ mice), which was further pronounced in the same strain at the 9-month time point, and even more prominent in *Pkd1*^RC/RC^ kidneys obtained from the strains with more rapidly progressive PKD (129S6/SvEvTac or BALB/cJ, **Figure 1C, D**). Importantly, as we observed for PD-1^+^/CD8^+^ kidney T cells, in mice with more severe and more rapidly progressive PKD (129S6/SvEvTac or BALB/cJ *Pkd1*^RC/RC^ mice), the proportion of PD-L1 positive CD64^+^ macrophages increased significantly as percent of CD64^+^ cells compared to wildtypes and with disease progression (**Supplemental Table 1**).

Beyond TAMs, tumor cells are another key cell type expressing PD-L1(44). Paralleling this finding, we found expression of PD-L1 to be significantly increased on kidney epithelial cells (APN^+^ or EpCAM^+^) of *Pkd1*^RC/RC^ mice compared to wildtype in all three strains (**Figure 1E, F**). That this increase did not consistently amplify as disease progressed can potentially be explained by an overall significant decrease of kidney epithelial cells as a proportion of live cells with progressive PKD; in advanced PKD, epithelial cell numbers are likely displaced by fibroblasts, immune cells etc. (**Supplemental Table 2**). Hence, it is important to note, that in line with our above reported findings, the proportion of kidney epithelial cells expressing PD-L1 in *Pkd1*^RC/RC^ kidneys increased significantly as disease advanced and compared to wildtypes (**Supplemental Table 1**). Together, these data show that expression of the PD-1|PD-L1 immune checkpoint proteins is significantly increased on relevant cells within PKD kidneys of *Pkd1*^RC/RC^ mice compared to wildtypes, suggesting induction of the pathway and an immunosuppressive CME.

### Epithelial PD-L1 is upregulated in human PKD

To evaluate whether PD-L1 expression is also upregulated in ADPKD human epithelial cells, we performed western blotting for PD-L1 in immortalized *PKD1*^+/+^ cells (renal cortical tubular epithelial [RCTE] cells, wildtype) and *PKD1*^-/’^cells (9-12 cells, ADPKD). We found minimal PD-L1 protein expression in RCTE cells. However, 9-12 cells had a significant increase in PD-L1 levels (∼7-fold increase) compared to RCTE cells (**Figure 1G**). Next, we looked for PD-L1 staining in human PKD end-stage kidney sections by immunohistochemistry (IHC). Compared to normal human kidney (NHK) tissue, we found that PD-L1 expression was substantially higher in kidney epithelial cells of ADPKD and Autosomal Recessive PKD (ARPKD) end-stage kidneys. This increase in expression was predominantly marked in the lining of cysts, but also in part within the cystic interstitium (**Figure 1H**). This data suggests that the immune checkpoint is also engaged in human PKD, mirroring our murine findings.

### Genetic loss of Pd-l1 or immune checkpoint blockade via a monoclonal PD-1 targeting antibody does not ameliorate cystic kidney disease in early- or adult-onset ADPKD

With the observed induction of the PD-1|PD-L1 pathway in our ADPKD1 model compared to wildtype and a seemingly intrinsic upregulation of PD-L1 in human ADPKD cells and cystic epithelia, we hypothesized that disruption of the PD-1|PD-L1 interaction would ameliorate cystic kidney disease in our orthologous model. To test this, we crossed both the adult-onset C57Bl/6J *Pkd1*^RC/RC^ mice with slowly progressive PKD and the adult-onset BALB/cJ *Pkd1*^RC/RC^ mice with rapidly progressive PKD to strain-matched *Pd-l1* (NCBI gene ID: *Cd274*) knockouts and compared PKD-associated phenotypes of *Pkd1*^RC/RC^;*Cd27*4^+/+^ to *Pkd1*^RC/RC^;*Cd27*4^-/-^ mice (**Figure 2A-D**). C57Bl/6J mice were aged till 9 months of age to assure we would not miss a possible impact on PKD phenotype at mild to moderate disease stages. BALB/cJ mice were euthanized at 3 months of age, as that timepoint represents the peak of cystic kidney disease severity after which cyst area and size regress due to cyst collapse and increased fibrotic burden(19, 41, 43). We assessed percent kidney weight/body weight (%KW/BW), cystic index (kidney cross-sectional area occupied by cysts), cyst size, cyst number, fibrotic index, and kidney function measured as blood urea nitrogen levels (BUN, **Figure 2B-D**, **Supplemental Table 3**). Surprisingly, we saw no significant difference in any of the measured parameters when comparing 9-month-old C57Bl/6J *Pkd1*^RC/RC^;*Cd274*^+/+^ to *Pkd1*^RC/RC^;*Cd274*^-/-^ mice or 3-month-old BALB/cJ *Pkd1*^RC/RC^;*Cd274*^+/+^ to *Pkd1*^RC/RC^;*Cd274*^-/-^ mice.

**Figure 2.**
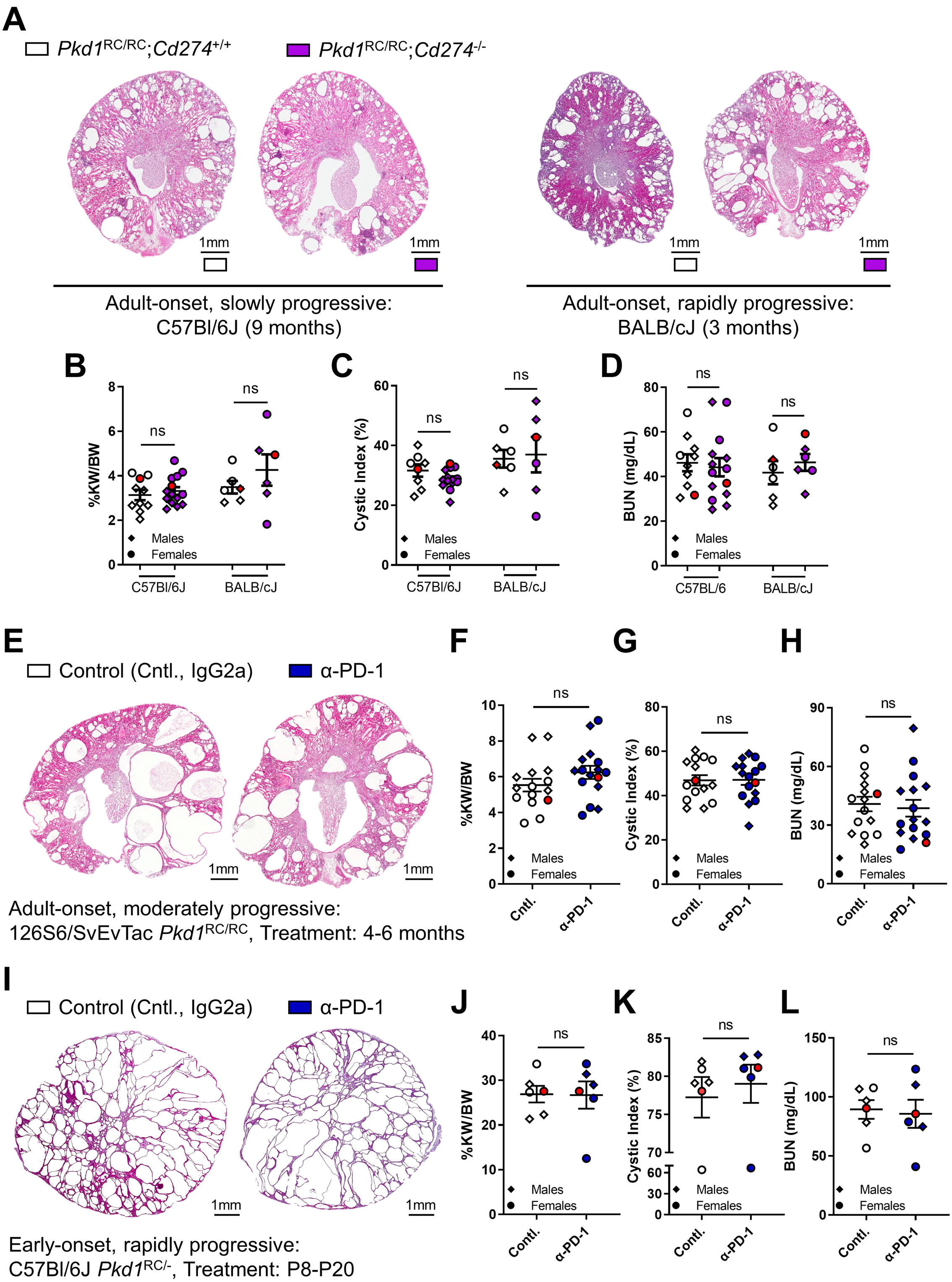
Genetic loss of *Pd-l1* or monoclonal α-PD-1 treatment does not impact polycystic kidney disease severity in mice with slowly or rapidly progressive ADPKD. (**A-D**) *Pkd1*^RC/RC^ mice were crossed with *Pd-l1*^-/-^ (NCBI gene ID: *Cd274*) mice to obtain *Pkd1*^RC/RC^;*Cd274*^+/+^ (control group; white) and *Pkd1*^RC/RC^;*Cd274*^-/-^ (experimental group; purple) mice. In the C57Bl/6J strain, animals were euthanized at 9 months, in the BALB/cJ strain at 3 months of age. (**E-H**) 129S6/SvEvTac *Pkd1*^RC/RC^ were treated with 10mg/kg of α-PD-1 (blue) or IgG2a (control, cntl., white) twice weekly by intraperitoneal injection from 4-6 months of age. (**I-L**) C57Bl/6J *Pkd1*^RC/-^ mice were treated with 10mg/kg of α-PD-1 (blue) or IgG2a (control, cntl., white) every other day by subcutaneous injection from postnatal day (P) 8 until P20. (**A, E, I**) Representative H&E cross-sectional images of the kidneys. Scale bars = 1mm. (**B, F, J**) %kidney weight/body weight (%KW/BW). (**C, G, K**) cystic index as measured by kidney area occupied by cysts (%). (**D, H, L**) blood urea nitrogen (BUN) levels. There was no difference in any of the analyzed PKD parameters, suggesting that inhibition of the PD-1/PD-L1 immune checkpoint does not impact adult-onset or early-onset PKD progression (also refer to **Supplemental Table 3**). Data are presented as the mean ± standard error of the mean (SEM); single data points are depicted. Males: diamond; females: circle. Red data points indicate the animal shown in (**A, E, or I**). Non-parametric Mann-Whitney tests were performed. ns, non-significant. (**A-D**) N=10-14 for C57Bl/6J or N=6 for BALB/cJ mice per experimental group. (**E-H**) N=15-16 mice per group, (**I-L**) N=6 mice per group.

Given the unexpected negative results from the genetic ADPKD1/PD-L1 cross, we wanted to assure the robustness of our findings using a secondary approach – inhibition of the PD-1|PD-L1 pathway via monoclonal antibody therapy. This additional step was especially important as in the genetic models the ability to engage the PD-1|PD-L1 immune checkpoint is lost from the time point of conception, potentially triggering other compensatory mechanisms (e.g., activation of other immunosuppressive pathways). Here, we chose to utilize an anti-PD-1 antibody for three reasons: [1] meta-analyses of anti-PD-1 vs. anti-PD-L1 treatment showed improved efficacy in non-small cell lung cancer (NSCLC) patients receiving anti-PD-1, [2] anti-PD-1 also allows for blockage of the PD-1/PD-L2 interaction, PD-L2 being a second ligand of PD-1, which when highly expressed in solid tumors has been shown to correlate with worse clinical outcome, and [3] anti-PD-1 inhibition complemented our prior-tested genetic approach by targeting the receptor versus the ligand(45, 46).

First, we treated 4-month-old 129S6/SvEvTac *Pkd1*^RC/RC^ mice for two months, twice weekly, by intraperitoneal (IP) injection with either 10mg/kg IgG2a (control) or 10mg/kg anti-PD-1 (α-PD-1). Using the 129S6/SvEvTac *Pkd1*^RC/RC^ model allowed us to test the therapeutic efficacy of PD-1|PD-L1 inhibition in a second adult-onset model with more rapidly progressive disease, complementing the BALB/cJ genetic studies. Second, we treated C57Bl/6J *Pkd1*^RC/-^ mice with either 10mg/kg IgG2a (control) or 10mg/kg anti-PD-1 (α-PD-1) injected subcutaneously every other day from postnatal day (P) 8 until P20. C57Bl/6J *Pkd1*^RC/-^ mice present with embryonic kidney cysts that rapidly advance, resulting in death by ∼P25(38, 41). Hence, this model allowed us to test the therapeutic efficacy of immune checkpoint blockade in an early-onset, rapidly progressive ADPKD model. In both instances we analyzed %KW/BW, cystic index, cyst size, cyst number, fibrotic index, and BUN after completion of the monoclonal therapy trial. Again, we saw no difference in any of the analyzed PKD parameters (**Figure 2E-L**, **Supplemental Table 3**). Taken together, these data indicate that genetic loss of *Pd-l1* or anti-PD-1 intervention has no effect on kidney cyst growth in our models, despite our data supporting a role for the PD-1|PD-L1 pathway in ADPKD.

### A combination strategy to augment immune checkpoint inhibitor efficacy led to decreased PKD severity in the BALB/cJ Pkd1^RC/RC^ model

There are greater than seven different immune checkpoints of T cell activation that have been targeted therapeutically in clinical cancer trials(47, 48). Among these, CTLA-4 and PD-1 have been found to be the most robust in reactivating CD8^+^ T cells. Indeed, ipilimumab, which blocks CTLA-4, was the first immune checkpoint antibody approved by the FDA for treatment of melanoma(49, 50). CTLA-4 functions by binding to CD80 and CD86, the ligand for the T cell co-stimulatory receptor CD28. CTLA-4 outcompetes CD28 to bind with CD80/86 and thereby dampens the release of pro-effector and cytotoxic cytokines in effector T cells(39). Multiple clinical trials were conducted to test the safety and efficacy of combining anti-PD-1 with anti-CTLA-4 which showed a remarkable increase in response rate and median survival time in multiple cancers including melanoma, RCC, NSCLC, and hepatocellular carcinoma, resulting in approval of the ipilimumab and nivolumab (anti-PD-1) combination for their treatment(39, 40, 51, 52). Of critical importance, combination treatment showed increased response rates in patients that did not benefit from monotherapy(39, 40).

Using scRNAseq, we recently published that both *Ctla4* and *Pdcd1*, encoding PD-1, are expressed in CD8^+^ T cells of mice lacking *Ift88* in the kidney and presenting with slowly progressive PKD(53). Based on these data, and the supportive rationale to combine anti-PD-1 and anti-CTLA-4 from the cancer field, we decided to test this approach in PKD. We chose the BALB/cJ *Pkd1*^RC/RC^ model for this experiment, as it would allow us to test anti-PD-1 monotherapy in an additional model without repeating our above experiments using the 129S6/SvEvTac *Pkd1*^RC/RC^ model. 1-month-old BALB/cJ *Pkd1*^RC/RC^ mice were treated twice a week for eight weeks by IP injection with either 10mg/kg anti-PD-1, 10mg/kg anti-CTLA-4, the combination of anti-PD-1 and anti-CTLA-4, or control IgG (10mg/kg IgG2a plus 10mg/kg IgG2b control). At 3 months of age the animals were euthanized and the same PKD parameters were analyzed as previously described. Like the outcomes in the 129S6/SvEvTac *Pkd1*^RC/RC^ model, monotherapy with anti-PD-1 did not impact PKD severity (**Figure 3, α-PD-1 [blue]**). Similarly, monotherapy with anti-CTLA-4 did not slow cystic kidney disease growth in this model (**Figure 3, α-CTLA-4 [green]**). Remarkably though, when we compared the animals that received both α-PD-1 and α-CTLA-4 to control, we found a significant reduction in nearly all the PKD parameters analyzed: %KW/BW, cystic index, fibrotic index, and BUN (**Figure 3**). Of note, kidney cyst number did not significantly change, but cyst size showed a trend towards smaller cysts in the mice that received both α-PD-1 and α-CTLA-4 suggesting that combination treatment predominantly impacted cyst expansion, and not cyst number. No health concerns indicative of adverse events were noted in the dually treated mice. Taken together, these data indicate that while monotherapy inhibiting immune checkpoint signaling does not slow PKD progression, suggesting some redundancy in the system, dual immune checkpoint inhibition significantly impacts kidney cyst growth in our orthologous ADPKD model.

**Figure 3.**
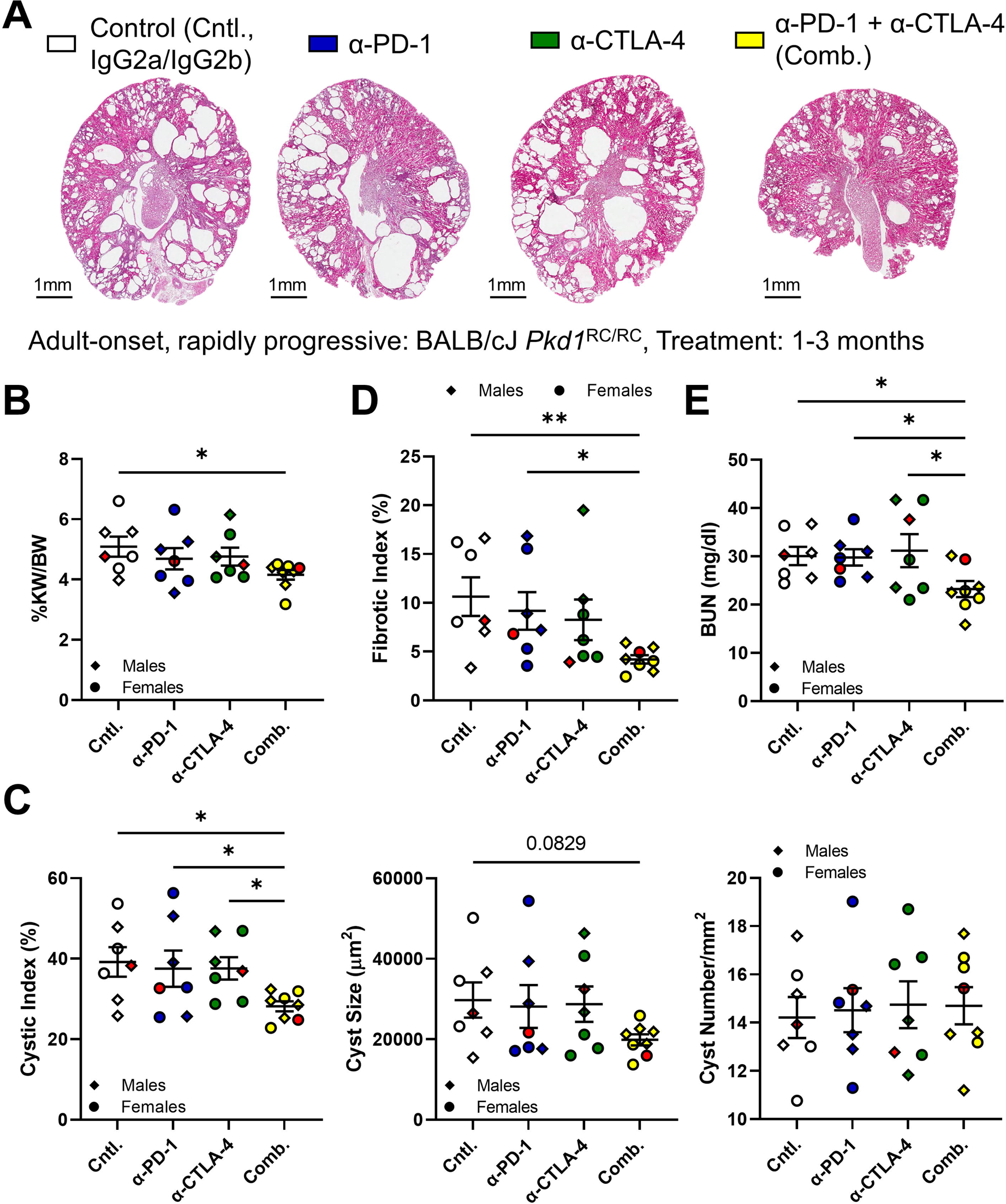
Combination immune checkpoint inhibition reduces cystic kidney disease in *Pkd1*^RC/RC^ mice. BALB/cJ *Pkd1*^RC/RC^ mice were treated from 1-3 months of age. Experimental groups: control (cntl., white, IgG2a plus IgG2b), α-PD-1 (blue, plus IgG2b), α-CTLA-4 (green, plus IgG2a), and combination (comb., yellow, α-PD-1 plus α-CTLA-4). (**A**) Representative H&E cross-sectional images of the kidneys. Scale bars = 1mm. (**B**) %kidney weight/body weight (%KW/BW). (**C**) cystic index as measured by kidney area occupied by cysts (%), average cyst size, and average cyst number normalized by tissue area. (**D**) fibrotic index (**E**) blood urea nitrogen (BUN) levels. Data are presented as the mean ± standard error of the mean (SEM); single data points are depicted. Males: diamond; females: circle. Red data points indicate the animal shown in (**A**). Kruskal-Wallis one-way analysis of variance with multiple comparison follow-up by controlling for false discovery rate (FDR, Benjamini, Krieger, Yekutieli) were performed. *p<0.05. N=7-8 mice per group.

### Combination immune checkpoint blockade leads to a more robust increase in kidney CD8^+^ T cell numbers/activation and reduced regulatory CD4^+^ T cell numbers compared to monotherapy alone

To better understand why monotherapy with either anti-PD-1 or anti-CTLA-4 did not alleviate PKD severity in our model but combination therapy did, we performed flow cytometry on kidneys at the end of the study to evaluate kidney CD8^+^ T cell numbers and activation status as defined by co-expression of CD44^+^/CD69^+^(19). In all treatment arms, we did not see a difference in overall kidney immune cell numbers (CD45^+^) compared to control, indicating that these treatments did not globally alter immune cell infiltration into the kidney (**Figure 4A**). Both anti-PD-1 as well as anti-CTLA-4 resulted in an increase in kidney CD8^+^ T cell numbers; importantly, there appeared to be an additive effect in animals that received the combination treatment (**Figure 4B**). When evaluating kidney CD8^+^ T cell activation status, we found very few CD8^+^ T cells to be expressing CD44^+^/CD69^+^ in the untreated *Pkd1*^RC/RC^ mice. Given this low frequency of effector CD8^+^ T cells may explain why CD8^+^ T cell loss worsens PKD, as we previously published(19). Both anti-PD-1 and anti-CTLA-4 treatment resulted in a moderate increase of CD8^+^, CD44^+^/CD69^+^ T cells in the kidney compared to control, ∼32% and ∼80% increase, respectively. Again, dual treatment with anti-PD-1 and anti-CTLA-4 increased the numbers of activated CD8^+^ T cells by ∼154% compared to control, which is an additional one-fold increase above the levels achieved by monotherapy (**Figure 4C**). Similarly, we found a three-fold increase in proliferating kidney CD8^+^ T cells (Ki67^+^) in animals receiving dual therapy compared to controls, while this increase was only 2-fold in the monotherapy arms (**Figure 4D**). Together, these data show that although there was no impact on PKD phenotype, both anti-PD-1 and anti-CTLA-4 monotherapies did result in an increase in numbers and activation status of CD8^+^ T cells. The fact that PKD progression was only slowed in animals receiving combination therapy suggests that there may be a threshold level of CD8^+^ T cell number/function needed to halt cyst growth.

**Figure 4.**
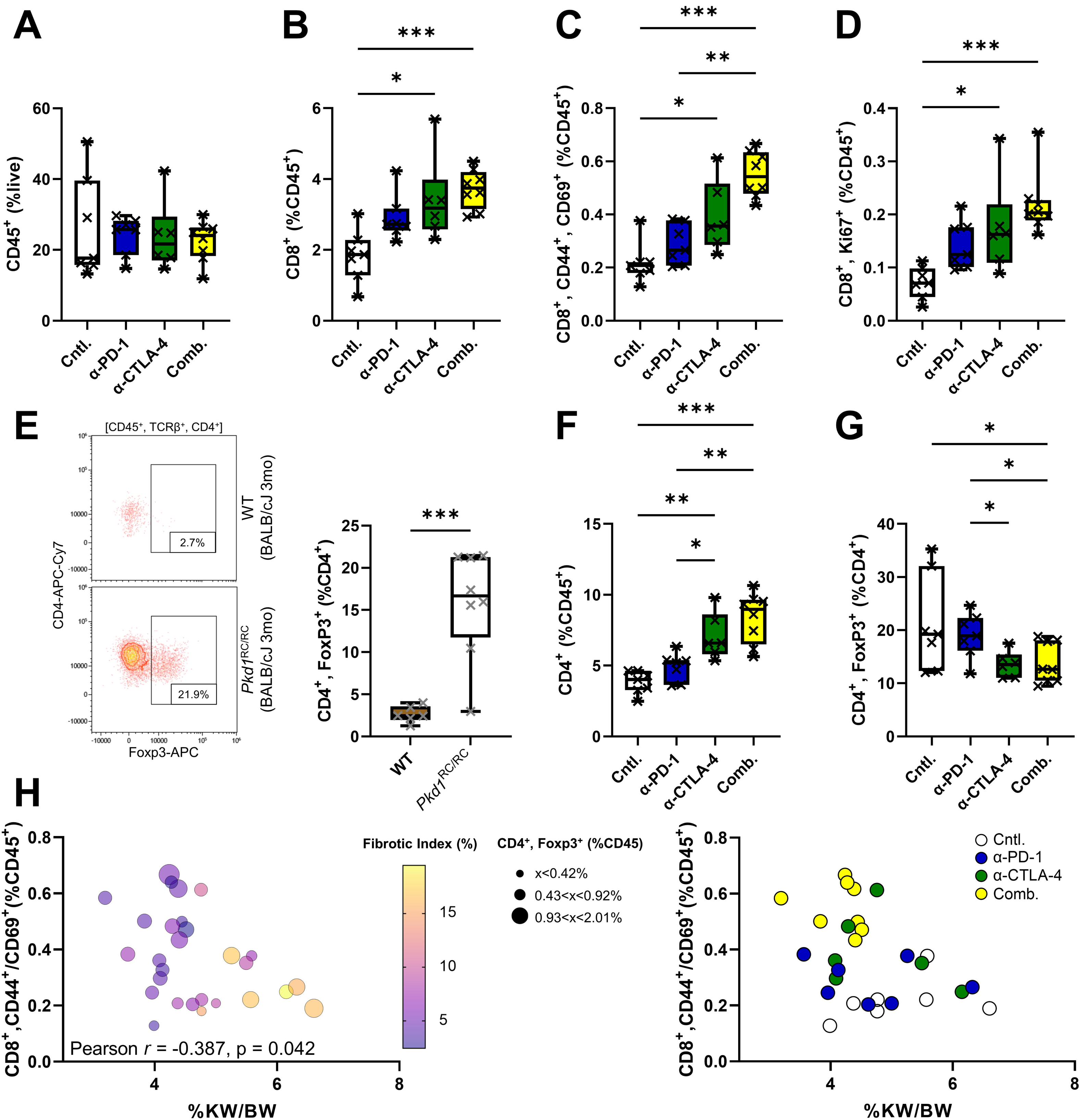
Treatment with α-PD-1 plus α-CTLA-4 results in rebalancing of the adaptive immunity within the cystic immune microenvironment. (**A-G, except E**) Flow cytometry of kidney single cell suspensions obtained at the end of treatment with either α-PD-1 (blue), α-CTLA-4 (green), or the combination (Comb., yellow) in BALB/cJ *Pkd1*^RC/RC^ mice compared to control (Cntl., white). (**A**) number of kidney immune cells (CD45^+^) as percent live. (**B**) number of kidney CD8^+^ T cells as percent CD45^+^. (**C, D**) number of kidney CD8^+^ T cells expressing CD44 and CD69, markers of T cell activation (**C**) or Ki67, marker of proliferation (**D**) as percent CD45^+^. Monotherapy with either α-PD-1 or α-CTLA-4 increased CD8^+^ T cell numbers, activation, and proliferation; however, the effect was amplified by combination treatment. (**F**) numbers of kidney CD4^+^ T cells as percent CD45^+^. (**G**) number of kidney CD4^+^ regulatory T cell numbers (T_Regs_, FoxP3^+^) as percent CD4^+^. While overall numbers of CD4^+^ T cell increased with each monotherapy, with additive effects seen in the combination treatment, numbers of T_Regs_ decreased within the treatment groups receiving α-CTLA-4 compared to control. (**E**) Analysis of kidney T_Reg_ numbers in 3-month-old BALB/cJ *Pkd1*^RC/RC^ mice (white) compared to wildtypes (brown) using flow cytometry. Representative flow diagrams (left), quantification (right). There is a highly significant increase in T_Reg_ numbers in ADPKD mice compared to wildtypes. (**H**) Correlation analyses of CD8^+^ T cell activation status and %KW/BW using all animals included in the immune checkpoint blockade combination trial. Left: each animal (individual data point) is color coded by the severity of fibrotic burden and sized by the number of kidney T_Regs_. There is a distinct correlation between animals with the least severe PKD (small %KW/BW and small fibrotic index) and high levels of activated CD8^+^ T cells combined with low levels of T_Regs_. Right: the same correlation plot as on the left but data points are color coded by treatment groups. See also **Supplemental Table 4**. (**A-G**) Data are presented as box plot with whiskers of 10-90th percentile; single data points are depicted. (**A-D, F, G**) Kruskal-Wallis one-way analysis of variance with multiple comparison follow-up by controlling for false discovery rate (FDR, Benjamini, Krieger, Yekutieli) were performed. (**E**) Non-parametric Mann-Whitney tests were performed. (**H**) Pearson correlation was performed using two-tailed multivariate analyses. *p<0.05, **p<0.01, ***p<0.001. (**A-D, F-H**) N=7-8 mice per group. (**E**) N=6-8 mice per genotype. Non-significant pair-wise comparisons are not shown.

In cancer, it is well established that T_Regs_ suppress anticancer immunity via a multitude of mechanisms(54). Interestingly, T_Regs_ express CTLA-4 as part of their function to maintain immune tolerance. Correlatively, recent cancer studies have shown that treatment with anti-CTLA-4 results in a reduction of intra-tumoral T_Regs_, representing a noncanonical mechanism that may contribute to therapeutic efficacy(55–57). The relevance of T_Regs_ to PKD progression has not been studied, but we recently found an enrichment of this population in PKD kidneys of an adult-onset PKD model caused by inducible kidney-specific *Ift88* loss versus controls(53). We confirmed that kidney T_Reg_ numbers are significantly increased in BALB/cJ *Pkd1*^RC/RC^ mice at 3 months of age compared to age, strain, gender matched wildtypes; the 3-month time point representing the endpoint of our combination therapy study (**Figure 4E**). Based on these findings, we analyzed the CD4^+^ T cell population in our anti-PD-1/anti-CTLA-4 study. Interestingly, kidney CD4^+^ T cell numbers increased as percent of all immune cells (CD45^+^), similarly to CD8^+^ T cells (**Figure 4F**). However, it is important to note that CD4^+^ T cells are comprised of multiple subtypes that have opposing function in immune regulation(58). Focusing on T_Regs_ as percent of CD4^+^ T cells, we found a 54% decrease in kidney T_Reg_ numbers in animals treated with anti-CTLA-4 or the combination compared to control, paralleling the cancer findings, and suggesting that this effect may have contributed to the efficacy of the combination treatment (**Figure 4G**).

To better understand the relevance of our observed adaptive immune cell changes to PKD severity, we correlated these flow cytometry parameters to %KW/BW, cystic index, and fibrotic index. Intriguingly, the number of CD8^+^ T cells that are activated, as characterized by CD44^+^/CD69^+^, were significantly inversely correlated with %KW/BW, cystic index, and fibrotic index (**Figure 4H**, **Supplemental Table 4**). Neither CD8^+^ nor CD4^+^ T cell numbers, proliferating CD8^+^ T cells (Ki67^+^), or T_Regs_ alone significantly correlated with these parameters (**Supplemental Table 4**). Combining the activation status of CD8^+^ T cells with the proliferative status modestly enhanced the correlation but the strongest correlation with %KW/BW was achieved when accounting for CD8^+^ T cell activation/proliferation status and T_Reg_ numbers, with CD8^+^ activation/proliferation status as a positive variable and T_Reg_ number as a negative variable (**Supplemental Table 4**). Together, these data suggest that the observed therapeutic efficacy with dual immune checkpoint inhibitors is likely to be contributed by both reactivation of the CD8^+^ T cell effector compartment and alleviation of immune suppressive mechanisms conferred by T_Regs_, resulting in a net rebalancing of the adaptive immunity within the CME.

## DISCUSSION

In our previous work we showed that the number of kidney CD8^+^ T cells increases with PKD severity and that depletion of CD8^+^ T cells worsens disease outcome(19). This suggests that this population of adaptive immune cells functions in halting cyst growth. It is known that CD8^+^ T cells lose functionality through engagement of immune checkpoints. Here, we explored the relevance of immune checkpoints, especially PD-1|PD-L1, to kidney cyst growth using different orthologous ADPKD1 models with varying rates of PKD progression. Using flow cytometry of kidney single cell suspensions, we found significant upregulation of PD-1 on CD8^+^ T cells and PD-L1 on kidney epithelial cells and macrophages. Upregulated expression of PD-L1 was also confirmed in ADPKD cell lines and patient kidney samples versus controls (**Figure 1**). This suggests that CD8^+^ T cells may be actively inhibited by this pathway within the CME and likely fail to execute their proposed anti-cystogenic function. This finding is in line with our recently published findings using scRNAseq which showed an enrichment of CD8^+^ T cells that express the PD-1 transcript in slowly progressive, inducible PKD mice lacking kidney *Ift88* compared to controls(53). Following the cancer paradigm, upregulation of PD-L1 on kidney macrophages and epithelial cells implies an active commitment to drive immunosuppression, which would allow for the cystic epithelium to escape immune-mediated killing. Although, PD-L1 upregulation could also suggest that the epithelium is protecting itself from immune-mediated injury, as seen in the case of lupus nephritis, or that it is simply concomitant of epithelial cell remodeling.

We confirmed the induction of the PD-1|PD-L1 immune checkpoint pathway in *Pkd1*^RC/RC^ mice inbred into three different strains (C57Bl/6J, BALB/cJ, and 129S6/SvEvTac, **Figure 1**). This demonstrates that the immune checkpoint is activated with progressive PKD independent of the rate of cyst growth – as known BALB/cJ and 129S6/SvEvTac *Pkd1*^RC/RC^ mice present with much more rapidly progressive adult-onset PKD than C57Bl/6J *Pkd1*^RC/RC^ mice. Further, it provides evidence that inhibition of kidney CD8^+^ T cell function is characteristic of PKD independent of the immune cell composition within the kidney, which differs significantly in regards to immune cell types and numbers in different mouse strains as has been shown by us in the case of kidney CD4^+^ to CD8^+^ T cell ratio(19, 59, 60). Hence, our data proposes that upregulation of PD-1|PD-L1 within the CME is a robust finding independent of the rate of PKD progression or mouse strain and suggests that immunosuppression is a critical feature of the PKD kidney. While there has been no direct evidence that immunosuppression may be important to cyst growth in PKD, several lines of evidence have hinted towards it. For example, ample data demonstrate that kidney macrophages in PKD models have an M2-like phenotype that mimics that of TAMs(10, 18). This includes that PKD macrophages express characteristic markers such as CD206, as well as upregulate the expression of ARG1, IL-10, CCR2, and CSF1R(14, 15, 61, 62). In line with these findings, depletion of macrophages as well as inhibition of CCR2 or CSF1R have been shown to slow cyst growth in inducible, early-onset or inducible, late-onset PKD models(10, 14, 15, 18). Beyond macrophages, our previous publication using inducible kidney-specific *Ift88* knockout mice, as well as our data here (**Figure 4E**) highlight an enrichment of T_Regs_ in PKD kidneys(53). T_Regs_, just like TAMs, are known drivers of immunosuppression in cancer(63).

Given our data that the PD-1|PD-L1 immune checkpoint is activated in our models, we tested the functional relevance of the pathway to PKD progression. Surprisingly, inhibiting the immune checkpoint both genetically and pharmacologically had no impact on disease severity (**Figure 2**). We confirmed these negative findings extensively focusing both on the receptor (PD-1) as well as the ligand (PD-L1) and testing the importance of the pathway to PKD pathogenesis in early-onset (*Pkd1*^RC/-^) as well as adult-onset (*Pkd1*^RC/RC^) ADPKD models. These results led us to conclude that PD-1|PD-L1 is likely not the only immune checkpoint engaged in the CME. Activation of additional immune checkpoint pathways would provide the opportunity to compensate our inhibitory interventions focused solely on PD-1|PD-L1 and explain why cyst growth was not retarded. Indeed, utilizing a combination therapy approach of anti-PD-1 and anti-CTLA-4 in BALB/cJ *Pkd1*^RC/RC^ mice slowed PKD progression compared to either monotherapy arm or control (**Figure 3**). We focused on CTLA-4 because we knew the gene is expressed in CD8^+^ T cells of inducible, kidney specific *Ift88* knockout mice presenting with adult-onset PKD, and because the monoclonal antibody targeting this receptor has proven efficacious in multiple cancers leading to FDA approval of the drug as monotherapy and combination therapy with anti-PD-1(39, 40, 51–53).

In line with cancer findings, kidney flow cytometry analyses of animals treated with both ICIs showed a clear reactivation of the effector T cell compartment, with increase CD8^+^ T cell numbers, activation, and proliferation. In addition, we further observed changes in the immunosuppressive T_Reg_ population associated with anti-CTLA-4 treatment (**Figure 4**). While some of these parameters were also observed in the monotherapy arms, correlation analyses with PKD phenotypes suggests that the combination of the analyzed adaptive immunity changes yielded the largest impact on slowing PDK progression. Seeing a therapeutic effect in slowing kidney cyst growth with the combined immune checkpoint blockade and an associated remodeling of the adaptive immunity not only provides, for the first time, functional evidence that immune checkpoint activation is a pathogenic driver of PKD but also re-emphasized the role of adaptive immune cells, specifically CD8^+^ T cells, in modulating PKD progression. Though, the therapeutic effect of dual immune checkpoint blockade, while significant, was only moderate, signifying that immunosuppression is only one of many pathways driving kidney cyst growth and accentuating the potential to combine therapeutic approaches that target both epithelial cells and cells within the CME.

Some caution is warranted when contemplating ICIs for PKD. In cancer patients ICI use has been associated with a variety of autoimmune adverse events involving the skin, the gastrointestinal tract, the endocrine system, and, to a lesser extent, the kidney. Approximately 2-3% of cancer patients receiving ICI monotherapy and ∼5% receiving ICI combination therapy present with acute kidney injury (AKI)(64, 65). Speculatively, ICIs break self-tolerance and trigger an autoimmune response against self-antigens which leads to immune-mediated injury in the kidney. While AKI can be managed clinically, this side effect cannot be ignored in the case of PKD patients, where kidney injury has been proposed to accelerate cyst growth in inducible PKD models(66, 67). Hence, long-term use of ICIs, as needed for the treatment of ADPKD, is likely not translatable. However, several cancer studies suggest a durable response and progression-free survival after discontinuation of mono- or combination ICI therapy for 6-12 months depending on the type of cancer(68–70). The underlying mechanisms associated with this treatment-free survival remain unclear but likely correspond to a permanent remodeling of the TME associated with innate and adaptive immune memory. Consequently, an altered dosing regimen which does not require continual long-term treatment but instead initial challenges and rechallenges with ICIs may be feasible to translate to the ADPKD clinic. However, the efficacy or side effects of such protocols were not tested in our studies and are part of future work alongside with investigations focused on alternative pathways that induce immunosuppression and CD8^+^ T cell exhaustion, which may be targetable alone or in combination with ICIs or other epithelial-centric drugs such as tolvaptan.

Our study has limitations. For one, we heavily focused on the *Pkd1* p.R3277C model. While we utilized mice spanning the full spectrum of ADPKD severity seen in patients, from early-onset (C57Bl/6J *Pkd1*^RC/-^) to late-onset (C57Bl/6J *Pkd1*^RC/RC^) PKD, the underlying disease driver was always a germline modification impacting polycystin-1 dosage. It is plausible that the therapeutic outcome of intervening with PD-1|PD-L1 signaling may differ when using other models of PKD such as germline *Pkd2* or ciliopathy models, as well as inducible kidney specific *Pkd1* or *Pkd2* knockout models. It is important though to keep the relevance of the model to human disease in mind, which was our priority when choosing the *Pkd1* p.R3277C model for these studies. Not only does the CME composition differ between rapidly progressive and slowly progressive models of PKD, as we have previously highlighted, it is also conceivable that the activation status/functionality of kidney immune cells varies significantly between germline and inducible PKD models(19). In PKD models driven by germline mutations kidney immune cells become conditioned to cyst-induced kidney remodeling from early embryogenesis on, while in inducible models the kidney immune cells remain naïve to cyst growth-associated changes prior PKD induction, changing the timescale for immune responses drastically. Our study was also limited in the cell types analyzed that express PD-1|PD-L1. We focused our analyses on the most studied and best characterized cell types, CD8^+^ T cells for PD-1 and epithelial cells/macrophages for PD-L1. However, the expression spectrum of PD-1 on adaptive immune cells goes far beyond CD8^+^ T cells. The receptor is expressed on different CD4^+^ T cells such as T_Regs_, on B cells, and on natural killer T cells. In addition, PD-1 has been found to be expressed on antigen presenting cells (APCs) as well as tumor cells, which has been associated with ICI therapy resistance(71). PD-L1 is also expressed on APCs beyond macrophages, such as dendritic cells, as well as effector T cells, where its functional relevance has been less well defined. We also did not evaluate the expression of CTLA-4|CD80/CD86 in a naïve setting or in the setting of PD-1|PD-L1 inhibition in our orthologous models. It is possible, that anti-PD-1 monotherapy showed no therapeutic efficacy due to activation of other immune checkpoints upon PD-1|PD-L1 blockade. Hence, more detailed investigations on cell-type specific expression of these immune checkpoint proteins and others, such as TIM-3 and LAG-3, and their relevance to PKD are still needed. Further, it would be important to prove that beyond activation and proliferation, CD8^+^ T cells also gained cytolytic activity and production of effector cytokines upon immune checkpoint blockade. Similarly, it would be interesting to identify changes in effector CD4^+^ T cell populations such as T_h_1 or T_h_17s as well as innate immune cell populations upon immune checkpoint blockade, neither of which was part of our studies. Hence, while our study provides critical foundational insight that immunosuppression (i.e., immune checkpoint activation) is a characteristic of the cystic kidney that functions in modulating disease severity, much more research is needed in understanding the underlying mechanism driving this phenomenon and targeting it therapeutically.

## CONCISE METHODS

Full methods are available in the **Supplemental Material**.

### Murine Study Details, Experimental Models, and Genetic Crosses

Fully inbred, homozygous C57Bl/6J, 129S6/SvEvTac, and BALB/cJ *Pkd1*^RC/RC^ (Pkd1^tm1.1Pcha^) mice and inbred, heterozygous C57Bl/6J *Pkd1*^del2/+^ (Pkd1^tm1Shh^) mice were obtained from the Mayo Clinic in 2015 (Dr. Peter C. Harris)(19, 38, 41, 42, 72). Fully inbred, homozygous C57Bl/6J and BALB/cJ *Cd274* knockout (Cd274^tm1Lpc^) mice were obtained in 2017 from Dr. Haidong Dong (Mayo Clinic) (73).

C57Bl/6J *Pkd1*^RC/-^ mice were obtained by crossing C57Bl/6J *Pkd1*^RC/RC^ and C57Bl/6J *Pkd1*^del2/+^ mice. C57Bl/6J or BALB/cJ *Pkd1*^RC/RC^;*Cd274*^+/+^ and *Pkd1*^RC/RC^;*Cd274*^-/-^ animals were obtained by crossing strain-matched F1 *Pkd1*^RC/RC^;*Cd274*^+/-^ pups. Housing and genotyping information are outlined in the **Supplemental Material**.

For all studies, both sexes, males and females, were utilized. Statistical analyses revealed no difference between males and females regarding evaluated PKD phenotypes; hence, both sexes were combined for all analyses.

### Human Samples

De-identified human ADPKD/Autosomal Recessive PKD (ARPKD) and normal human kidney (NHK) FFPE sections were obtained through an MTA with the Kansas PKD Research and Translational Core Center (P30 DK106912) at the University of Kansas Medical Center. The PKD samples are end-stage kidney samples which were collected post-transplant.

### Cell Culture

All cell lines have been previously described(74); renal cortical tubular epithelial (RCTE) cells, *PKD1*^+/+^, and 9-12 cells, *PKD1*^-/-^. Cells were grown in Dulbecco’s modified Eagle’s medium/Ham’s F-12 50/50 mixed with L-glutamine and 15nM HEPES (DMEM/F12; *Corning*) supplemented with 10% fetal bovine serum (*Sigma-Aldrich*) and 1% penicillin-streptomycin (*Corning*) for less than 10 passages.

### Immunoblotting

RCTE and 9-12 cells were grown until ∼70-80% confluency and protein was isolated. A total of 30µg of protein were analyzed on a 10% sodium dodecyl-sulfate polyacrylamide gel, transferred to a polyvinylidene difluoride membrane, and incubated with primary antibody overnight. The membranes were exposed to ECL reagent (*Perkin Elmer*, #NEL104001EA) and developed using an X-ray film developer. Band density of each blot was quantified using ImageJ software. Antibodies: rabbit anti-PD-L1 (*Cell Signaling Technology*, #13684; 1:1,000), goat anti-rabbit-HRP (*Pierce*, #1858415; 1:5,000), mouse anti-β-actin (*Sigma-Aldrich,* #A5441; 1:10,000), goat anti-mouse-HRP (*Jackson ImmunoResearch Laboratories Inc*, #115-035-003;1:20,000).

### Immunohistochemistry

Human kidney ADPKD, ARPKD and NHK sections were stained following the VectaStain Elite ABC Universal Plus kit (*Vector Laboratories*, #PK-8200). Antibodies: Anti-PD-L1 (*Abcam*, #ab205921; 1:100).

### Histomorphometric and Kidney Function Analyses

Cystic index, cyst size, and cyst number were analyzed as previously published(19). Fibrotic area was analyzed from picrosirius red stained kidney sections visualized using an Olympus BX41 microscope with a linear polarizer as previously published(19).

Blood urea nitrogen (BUN) levels were analyzed following the manufacturer’s protocol (QuantiChrom Urea Assay Kit, #501079333, *BioAssay Systems*). Samples were analyzed in duplicates.

### Immunodepletion Experiments

#### Anti-PD-1 studies

Four-month-old 129S6/SvEvTac *Pkd1*^RC/RC^ mice were treated twice a week for eight weeks by intraperitoneal (IP) injection with 10mg/kg anti-PD-1 blocking antibody (clone RMP1-14; *Bio X Cell*) or 10mg/kg IgG2a control (clone 2A3; *Bio X Cell*). C57Bl/6J *Pkd1*^RC/-^ mice were treated every other day by subcutaneous injection with 10mg/kg anti-PD-1 blocking antibody or 10 mg/kg IgG2a control starting at postnatal day (P) 8 until P20.

#### Anti-PD-1/Anti-CTLA4 study

One-month-old BALB/cJ *Pkd1^RC/RC^* mice were treated twice a week for eight weeks by IP injection with 10 mg/kg anti-PD-1 blocking antibody 10 mg/kg anti-CTLA-4 blocking antibody (clone 9D9; *Bio X Cell*), the combination of anti-PD-1 and anti-CTLA-4, or control IgG (10 mg/kg IgG2a and 10 mg/kg IgG2b control [clone MPC-11, *Bio X Cell*]). Animals with single blockade of PD-1 or CTLA-4 also received the respective other control antibody.

### Single Cell Suspension & Flow Cytometry

Single cell suspensions of the dissected kidneys were prepared as previously described(19). In short, tissue was mechanically dissociated and digested in DMEM/F12 media (*Corning*) containing Liberase TL (2mg/mL, *Sigma-Aldrich*) and DNase I (20K U/mL, *Sigma-Aldrich*) for 30min at 37°C, then passed through a 100µm and 70µm filter, as well as cleared of red blood cells using red blood bell lysis buffer (0.015M NH_4_Cl, 10mM KHCO_3_, 0.1mM Na_2_EDTA, pH 7.2).

Staining of single cell suspension and flow cytometry protocol were described previously(19). Each kidney cell suspension was split in half and stained with two different panels (see below). The single cell suspension was blocked in anti-mouse CD16/CD32 (clone 93; *eBioscience*) for 15min, followed by viability staining (LIVE/DEAD Fixable Aqua Dead Cell Stain Kit, *Invitrogen*) for 15min and conjugated surface antibody staining for 30min. Intracellular markers were stained for according to the Foxp3/Transcription Factor Staining Buffer Set (*eBioscience*, #00-5523-00).

#### T cell Panel

CD44-FITC (clone IM7; 1:100; *eBioscience*), PD-1-PE (clone J43; 1:200; *eBioscience*), CD45-PE/CF594 (clone 30-F11; 1:100; *Fisher Scientific*), TCRβ-PE/Cy5 (clone H57-597; 1:200; *Affymetrix*), Ki- 67-APC (clone SolA15; 1:400; *eBioscience*) or FoxP3-APC (clone FJK-16s; 1:200; *eBioscience*), CD69-PE-Cy7 (clone H1.2F3; 1:100; *eBioscience*), CD8-AF700 (clone 53-6.7; 1:100; *eBioscience*), CD4-APC/Cy7 (clone GK1.5; 1:200; *BioLegend*), FoxP3-eFluor450 (clone FJK-16s; 1:200; *eBioscience*)

#### Epithelial/macrophage panel

CD45-FITC (clone 30-F11; 1:100; *eBioscience*), CD11c-PE (clone HL3; 1:100; *BD Pharmingen*), CD19-PE/AF594 (clone 1D3; 1:100; *BD Horizen*), Ly6G-PercP/Cy5.5 (clone 1A8; 1:100; *Biolegend*), PD-L1-PE/Cy7 (clone MIH5; 1:100; *eBioscience*), CD64-AF647 (clone ; 1:100), APN-AF700 (clone ER-BMDM1; 1:100; *Novus Biologicals*), EpCAM-APC/eFluor780 (clone G8.8; 1:100; *Invitrogen*).

Following staining, cells were ran on the Gallios Flow Cytometer Machine (*Beckman Coulter*) and analyzed using Kaluza Analysis v2.1 software (*Beckman Coulter*). The analyses workflow plus gating strategies are outlined in the **Supplemental Material**.

### Statistics

All analyses were performed using JMP Pro 16.1 (*SAS*) or PRISM 9.2.0 (*Graphpad Software*). Data are presented as the mean ± standard error of the mean (SEM) or box plot with whiskers of 10-90th percentile; single data points are depicted. Pairwise comparisons were performed using a Mann-Whitney test and multi-group comparisons were performed using Kruskal-Wallis test with multiple comparison follow-up by controlling for FDR (Benjamini, Krieger, Yekutieli). The Pearson correlation coefficient was computed using two-tailed multivariate analyses with a confidence interval of 95%. P-values are denoted by *(p<0.05), **(p<0.01), ***(p<0.001), and ****(p<0.0001).

### Study approval

All experimental procedures were performed in an AAALAC-accredited facility in accordance with the *Guide for the Care and Use of Laboratory Animals*(75) and approved by the University of Colorado Anschutz Medical Campus Institutional Animal Care and Use Committee (protocols #33, 301, 685).

## AUTHOR CONTRIBUTION

Conceived and designed research: E.K.K., E.T.C., R.A.N., K.H.; Performed experiments: E.K.K., D.T.N., K.H.M., C.D.B., A.S.L., S.B.F., K.H.; Analyzed data: E.K.K., D.T.N., K.H.; Interpreted results of experiments: E.K.K., D.T.N., E.T.C., K.A.Z., R.A.N., K.H.; Prepared figures: E.K.K., K.H.; Drafted manuscript: E.K.K., K.H.; Edited and revised manuscript: E.K.K., D.T.N., A.S.L., S.B.F., B.Y.G., M.B.C., E.T.C., K.A.Z.,

## ACKNOWLEDMENTS

Support was provided by the NIH NIDDK K01 DK119375 (KAZ), K01 DK114164 (KH), NIH NIDDK K01 DK114164-03S1 (KH), University of Colorado, Anschutz Medical Campus Center for Fibrosis Research and Translation Pilot & Feasibility Grant (RAN), NIH NIDDK T32 5T32DK007135 (EKK), PKD Foundation Research Grant 216G18a (KH), and the Zell Family Foundation (RAN, KH).

