## Supplemental Material for "Immune Checkpoint Activity Regulates Polycystic Kidney Disease Progression"

[+]Contributed equally to this work.

***Corresponding Author:**

Dr. Katharina Hopp

### SUPPLEMENTAL FULL METHODS

*Murine Study Details, Experimental Models, and Genetic Crosses*

All study animals were group-housed, separated by gender, with a maximum of five animals per cage, maintained on a standard diet (ENVIGO #2020), and had food plus water freely available. The housing facility is pathogen-free and was kept at a temperature of ~72ºF, humidity of ~37%, and a light cycle of 8pm off, 6am on.

Fully inbred, homozygous C57Bl/6J, 129S6/SvEvTac, and BALB/cJ *Pkd1*$^{RC/RC}$ mice (Pkd1$^{tm1.1Pcha}$) were obtained from the Mayo Clinic in 2015 (Dr. Peter C. Harris) with an approved material transfer agreement (MTA) and were maintained in homozygosity by the principal investigator (PI), Katharina Hopp (1-4). Homozygous *Pkd1*$^{RC/RC}$ mice were outcrossed every 10$^{th}$ generation to strain-matched wildtype mice obtained from *The Jackson Laboratory* (stock number 000664 [C57Bl/6J]; 000651 [BALB/cJ]) or *Taconic Biosciences* (model number 126SVE). Inbred, heterozygous C57Bl/6J *Pkd1*$^{del2/+}$ (Pkd1$^{tm1Shh}$) mice were obtained under the same MTA mentioned above and maintained in heterozygosity by crossing with C57Bl/6J wildtypes (*The Jackson Laboratory*, stock number 000664)(5).

Fully inbred, homozygous C57Bl/6J and BALB/cJ *Cd274* knockout mice (Cd274$^{tm1Lpc}$) were obtained in 2017 from Dr. Haidong Dong (Mayo Clinic) with the appropriate MTA in place(6). They were maintained in homozygosity and used for all experiments without outcrossing.

C57Bl/6J *Pkd1*$^{RC/-}$ mice were obtained by crossing C57Bl/6J *Pkd1*$^{RC/RC}$ and C57Bl/6J *Pkd1*$^{del2/+}$ mice. C57Bl/6J or BALB/cJ *Pkd1*$^{RC/RC}$;*Cd274*$^{+/+}$ and *Pkd1*$^{RC/RC}$;*Cd274*$^{-/-}$ animals were generated by crossing strain-matched F1 *Pkd1*$^{RC/RC}$;*Cd274*$^{+/-}$ pups obtained from four different *Pkd1*$^{RC/RC}$ and *Cd274*$^{-/-}$ parental crosses. In total, six different F1 crosses generated all F2 experimental animals (C57Bl/6J or BALB/cJ *Pkd1*$^{RC/RC}$;*Cd274*$^{+/+}$ and *Pkd1*$^{RC/RC}$;*Cd274*$^{-/-}$).

The genotypes of all experimental animals were PCR confirmed using DNA extracted from tail clips. The *Pkd1* p.R3277C genotyping protocol was performed as previously published(2). The *Pkd1* del2 genotyping was performed using a wildtype (5'-ACGCTGGGCAAAGGAACATC-3'; 5'- TGGGAACAGAGAGAGACAGTGGTC-3'; 380bp) and mutant (5'- CGACCACCAAGCGAAACATC-3'; 5'-GTCCGACATTGCTCCTGTGC-3'; 475 bp) PCR reaction. Both PCRs were run using the same conditions (94°C 3min; 35x 94°C 30sec, 59°C 30sec, 72°C 40sec; 72°C 10min; 4°C 10min). The *Cd274* genotyping was performed as multiplex PCR (5'-

AGAACGGGAGCTGGACCTGCTTGCGTTAG-3’; 5’-ATTGACTTTCAGCGTGATTCGCTTGTAG-3’; 5’-TCTATGGCTTCTGAGGCGGA-3’; wildtype: 250bp, mutant 160bp) using the following PCR conditions: 94°C 3min; 35x 94°C 30sec, 58°C 30sec, 72°C 50sec; 72°C 10min; 4°C 10min.

For all studies, both sexes, males, and females, were utilized. Statistical analyses revealed no difference between males and females regarding evaluated PKD phenotypes; hence, both sexes were combined for all analyses. All animals were aged as outlined in the results sections prior to euthanasia and PKD parameter analyses were performed as outlined below. The number of animals per study varied and is indicated in the respective results sections and figure legends.

*Human Samples*

De-identified human ADPKD/Autosomal Recessive PKD (ARPKD) and normal human kidney (NHK) FFPE sections were obtained through an MTA with the Kansas PKD Research and Translational Core Center (P30 DK106912) at the University of Kansas Medical Center. The PKD samples are end-stage kidney samples which were collected post-transplant.

*Cell Culture*

All cell lines have been previously described(7). Briefly, renal cortical tubular epithelial (RCTE) cells are *PKD1*$^{+/+}$ immortalized human renal cortical tubular epithelial cells, and 9-12 cells are *PKD1*$^{-/-}$ immortalized cells derived from human ADPKD cystic epithelium. Cells were grown in Dulbecco’s modified Eagle’s medium/Ham’s F-12 50/50 mixed with L-glutamine and 15nM HEPES (DMEM/F12; *Corning*) supplemented with 10% fetal bovine serum (*Sigma-Aldrich*) and 1% penicillin-streptomycin (*Corning*). Cells were grown in a humidified incubator with 5% $CO_2$ at 37°C. Cells were kept in culture for no longer than 6-8 weeks and up to 10 passages at maximum before a new vial was thawed.

*Immunoblotting*

RCTE and 9-12 cells were grown on tissue-culture treated 10cm plates until ~70-80% confluency, after which cells were harvested and lysed (0.5% Triton X-100, 50 mM β-glycerophosphate [pH 7.2], 0.1 mM $Na_3VO_4$,

2mM $MgCl_2$, 5 mM EDTA, plus protease inhibitor cocktail [*Sigma-Aldrich,* cOmplete™, #P8340]). Protein concentration was measured using protein assay dye reagent concentrate (*Bio-Rad*, #5000006): Bradford method. A total of 30μg of protein were loaded on a 10% sodium dodecyl-sulfate polyacrylamide gel and transferred onto a polyvinylidene difluoride membrane. Membranes were blocked in 5% bovine serum albumin in Tris-buffered saline with 0.1% Tween 20 (TBST) for 1 hour, incubated overnight at 4°C with primary antibodies, and for 1 hour with secondary antibody at room temperature. For probing of different antibodies, membranes were stripped using Restore Western Blot Stripper Buffer (*Thermo Scientific*) according to the manufacturer's instructions. The membranes were exposed to ECL reagent (*Perkin Elmer*, #NEL104001EA) and developed using an X-ray film developer. Band density of each blot was quantified using ImageJ software. Antibodies: rabbit anti-PD-L1 (*Cell Signaling Technology*, #13684; 1:1,000), goat anti-rabbit-HRP (*Pierce*, #1858415; 1:5,000), mouse anti-β-actin (*Sigma-Aldrich,* #A5441; 1:10,000), goat anti-mouse-HRP (*Jackson ImmunoResearch Laboratories Inc*, #115-035-003;1:20,000).

*Immunohistochemistry*

Human kidney ADPKD, ARPKD and NHK sections were stained following the VectaStain Elite ABC Universal Plus kit (*Vector Laboratories*, #PK-8200). Antigen unmasking was performed with slides emerged in Universal HIER antigen retrieval reagent (*Abcam*, #ab208572) at 100°C for 20min and 90°C for 20min. Endogenous peroxidase activity was quenched using BLOXALL blocking solution following the manufacturing protocol (*Vector Laboratories*, #SP-6000). Antibodies: Anti-PD-L1 (*Abcam*, #ab205921; 1:100). The negative control slide underwent the same protocol as described above with the exception that no primary antibody was added. The supplied secondary antibody from the VectaStain Elite ABC Universal Plus kit was used at the recommended dilution.

*Mouse Tissue Harvest*

For all animal experiments, mice were euthanized by isoflurane exposure and cervical dislocation, and the body weight of each animal was recorded. Following, terminal heparin blood was collected via cardiac puncture. For mice analyzed by flow cytometry, mice were perfused with 20mL ice cold PBS/heparin (80units/mL) via pressure perfusion. After perfusion or directly after blood collection, the kidneys and spleen were harvested and weighed. For animals analyzed by flow cytometry, the left kidney and half the spleen were used for the single cell suspension/flow cytometry and half of the right kidney plus the other half of the spleen were fixed in 4% paraformaldehyde for histological analyses. The other half of the right kidney was flash frozen. For all other animals, the right kidney and half of the spleen were fix in 4% paraformaldehyde for histological analyses and the left kidney plus the remainder of the spleen were flash frozen. The age of mice at euthanasia/tissue harvest varied depending on experiment (refer to results sections).

*Histomorphometric and Kidney Function Analyses*

Cystic index, cyst size, and cyst number were analyzed as previously published(4). In short, three kidney cross sections per animal (horizontal plane at pelvis, mid-superior and mid-inferior pole) were analyzed using a custom-built NIS-Elements AR v4.6 macro (*Nikon*) - a cyst was defined as having a minimum feret of 50µm. Fibrotic area was analyzed from picrosirius red stained kidney sections visualized using an Olympus BX41 microscope with a linear polarizer as previously published(4). Ten random cortical 10x images were analyzed per animal.

Blood urea nitrogen (BUN) levels were analyzed using the terminal blood collection and following the manufacturer's protocol (QuantiChrom Urea Assay Kit, #501079333, *BioAssay Systems*). Samples were analyzed in duplicates.

*Immunodepletion Experiments*

<u>Anti-PD-1 studies:</u>

Four-month-old 129S6/SvEvTac *Pkd1*$^{RC/RC}$ mice were treated twice a week for eight weeks by intraperitoneal (IP) injection with 10mg/kg anti-PD-1 blocking antibody (clone RMP1-14; *Bio X Cell*) or 10mg/kg IgG2a control (clone 2A3; *Bio X Cell*). After eight weeks of treatment, the animals were euthanized and evaluated as described above.

C57Bl/6J *Pkd1*$^{RC/-}$ mice were treated every other day by subcutaneous injection with 10mg/kg anti-PD-1 blocking antibody (clone RMP1-14; *Bio X Cell*) or 10 mg/kg IgG2a control (clone 2A3; *Bio X Cell*) starting at postnatal day (P) 8 until P20, at which point the animals were euthanized and evaluated as described above. For both studies, the depletion antibody and the control antibody were diluted in PBS.

Anti-PD-1/Anti-CTLA4 study:

One-month-old BALB/cJ *Pkd1*$^{RC/RC}$ mice were treated twice a week for eight weeks by IP injection with 10 mg/kg anti-PD-1 blocking antibody (clone RMP1-14; *Bio X Cell*), 10 mg/kg anti-CTLA-4 blocking antibody (clone 9D9; *Bio X Cell*), the combination of anti-PD-1 and anti-CTLA-4, or control IgG (10 mg/kg IgG2a [clone 2A3; *Bio X Cell*] and 10 mg/kg IgG2b control [clone MPC-11, *Bio X Cell*]). Animals with single blockade of PD-1 or CTLA-4 also received the respective other control antibody. Antibodies were diluted in InVivoPure pH 7.0 Dilution Buffer (*Bio X Cell*). After eight weeks of treatment, the animals were sacrificed and evaluated as described above. The doses given are standard for *in vivo* studies and have been extensively published on.

*Single Cell Suspension & Flow Cytometry*

Single cell suspensions of the dissected kidneys were prepared as previously described(4). In short, tissue was mechanically dissociated and digested in DMEM/F12 media (*Corning*) containing Liberase TL (2mg/mL, *Sigma-Aldrich*), and DNase I (20K U/mL, *Sigma-Aldrich*) for 30min at 37°C. The digestion mix was passed through a 100µm and 70µm filter, as well as cleared of red blood cells using red blood bell lysis buffer (0.015M $NH_4Cl$, 10mM $KHCO_3$, 0.1mM $Na_2EDTA$, pH 7.2). Cells were pelleted and prepped for flow cytometry. Staining of single cell suspension and flow cytometry protocol were described previously(4). Each kidney cell suspension was split in half and stained with two different panels (see below). The single cell suspension was blocked in anti-mouse CD16/CD32 (clone 93; *eBioscience*) for 15 min, followed by viability staining (LIVE/DEAD Fixable Aqua Dead Cell Stain Kit, *Invitrogen*) for 15min and conjugated surface antibody staining for 30min. For the T cell panel outlined below, the single cell suspension then underwent fixation, permeabilization, and intracellular antibody staining according to the Foxp3/Transcription Factor Staining Buffer Set (*eBioscience*, #00-5523-00). All incubations were done at 4°C.

T cell Panel: CD44-FITC (clone IM7; 1:100; *eBioscience*), PD-1-PE (clone J43; 1:200; *eBioscience*), CD45-PE/CF594 (clone 30-F11; 1:100; *Fisher Scientific*), TCRβ-PE/Cy5 (clone H57-597; 1:200; *Affymetrix*), Ki-67-APC (clone SolA15; 1:400; *eBioscience*) or FoxP3-APC (clone FJK-16s; 1:200; *eBioscience*), CD69-PE-Cy7 (clone H1.2F3; 1:100; *eBioscience*), CD8-AF700 (clone 53-6.7; 1:100; *eBioscience*), CD4-APC/Cy7 (clone GK1.5; 1:200; *BioLegend*), FoxP3-eFluor450 (clone FJK-16s; 1:200; *eBioscience*)

Epithelial/macrophage panel: CD45-FITC (clone 30-F11; 1:100; *eBioscience*), CD11c-PE (clone HL3; 1:100; *BD Pharmingen*), CD19-PE/AF594 (clone 1D3; 1:100; *BD Horizen*); Ly6G-PercP/Cy5.5 (clone 1A8; 1:100; *Biolegend*), PD-L1-PE/Cy7 (clone MIH5; 1:100; *eBioscience*), CD64-AF647 (clone ; 1:100), APN-AF700 (clone ER-BMDM1; 1:100; *Novus Biologicals*), EpCAM-APC/eFluor780 (clone G8.8; 1:100; *Invitrogen*).

Following staining cells were ran on the Gallios Flow Cytometer Machine (*Beckman Coulter*) and analyzed using Kaluza Analysis v2.1 software (*Beckman Coulter*). For analyses, compensation was performed using single antibody-stained beads (VersaComp, #B22804, *Beckman Coulter*) and single antibody-stained cells. The gating strategy for the T cell panel was performed as previously published (4) with the following addition: $CD8^+$ T cells were gated for staining positive for PD-1 or Ki-67. A fluorescence minus one (FMO) sample was used to set the negative gate.

Gating for PD-L1 positive epithelial cells and macrophages was done as follow: Live singlet cells were gated for CD45 positive or CD45 negative. CD45 negative cells were gated for being either $APN^+$ or $EpCAM^+$ using a Boolean "OR" gate (epithelial cells). Epithelial cells were then analyzed to be either positive or negative for PD-L1 staining. A FMO sample was used to set the negative gate (**Figure 1D**). Macrophages were defined by analyzing $CD45^+$ cells that were $CD19^-$ and $Ly6G^-$, to exclude B cells and neutrophils. Resulting cells were then gated for $CD64^+$ events, followed by analysis of PD-L1 positive expression using the same gate used for epithelial cells (**Figure 1C**).

*Statistics*

All analyses were performed using JMP Pro 16.1 (*SAS*) or PRISM 9.2.0 (*Graphpad Software*). Data are presented as the mean ± standard error of the mean (SEM) or box plot with whiskers of 10-90th percentile; single data points are depicted. Pairwise comparisons were performed using a Mann-Whitney test and multi-group comparisons were performed using Kruskal-Wallis test with multiple comparison follow-up by controlling for FDR (Benjamini, Krieger, Yekutieli). The Pearson correlation coefficient was computed using two-tailed multivariate analyses with a confidence interval of 95%. P-values are denoted by *(p<0.05), **(p<0.01), ***(p<0.001), and ****(p<0.0001).

*Study approval*

All experimental procedures were performed in an AAALAC-accredited facility in accordance with the *Guide for the Care and Use of Laboratory Animals*(8) and approved by the University of Colorado Anschutz Medical Campus Institutional Animal Care and Use Committee (protocols #33, 301, 685).

### SUPPLEMENTAL TABLES

#### Supplemental Table 1 | Population specific distribution of PD-1/PD-L1 expressing cells in mouse wildtype and ADPKD1 kidneys.

| | CD8$^+$, PD-1$^+$ (%CD8$^+$) | | | CD64$^+$, PD-L1$^+$ (%CD64$^+$) | | | EpCAM$^+$/APN$^+$, PD-L1$^+$ (%EpCAM$^+$/APN$^+$) | | |
|---|---|---|---|---|---|---|---|---|---|
| C57Bl/6J | WT | *Pkd1* $^{RC/RC}$ | *Statistics* | WT | *Pkd1* $^{RC/RC}$ | *Statistics* | WT | *Pkd1* $^{RC/RC}$ | *Statistics* |
| 3mo | 12.88±2.79 | 13.76±2.11 | ns | 1.30±0.10 | 1.26±0.23 | ns | 1.78±0.23 | 1.93±0.16 | ns |
| 6mo | 28.92±2.98 | 28.72±4.08 | ns/## | 1.82±0.32 | 2.13±0.30 | ns/# | 2.25±0.21 | 3.55±0.39 | **/### |
| 9mo | 37.54±3.56 | 31.80±2.97 | ns/###/ns | 4.48±0.66 | 4.67±0.55 | ns/####/^^ | 2.05±0.32 | 7.96±1.74 | ****/####/^ |
| 129S6/SvEVTac | WT | *Pkd1* $^{RC/RC}$ | *Statistics* | WT | *Pkd1* $^{RC/RC}$ | *Statistics* | WT | *Pkd1* $^{RC/RC}$ | *Statistics* |
| 3mo | 7.48±1.69 | 9.36±0.94 | ns | 0.94±0.18 | 2.28±0.24 | ** | 0.72±0.05 | 3.81±0.60 | *** |
| 6mo | 4.27±1.06 | 14.02±2.27 | **/ns | 1.29±0.17 | 2.70±0.32 | **/ns | 1.57±0.13 | 6.13±0.96 | ***/ns |
| 9mo | 1.56±0.74 | 7.05±1.23 | **/ns/^ | 0.91±0.12 | 1.92±0.20 | **/ns/ns | 1.04±0.13 | 4.62±0.59 | **/ns/ns |
| BALB/cJ | WT | *Pkd1* $^{RC/RC}$ | *Statistics* | WT | *Pkd1* $^{RC/RC}$ | *Statistics* | WT | *Pkd1* $^{RC/RC}$ | *Statistics* |
| 3mo | 0.35±0.12 | 9.28±0.98 | ** | 1.23±0.06 | 1.76±0.24 | ns | 0.78±0.07 | 1.92±0.24 | *** |
| 6mo | 2.65±0.32 | 26.92±3.49 | **/### | 1.19±0.13 | 2.25±0.41 | **/ns | 1.06±0.08 | 4.68±0.31 | **/### |
| 9mo | 8.59±4.07 | 25.15±2.17 | */###/ns | 4.45±1.93 | 3.36±0.35 | ns/##/^ | 2.25±0.65 | 7.07±0.69 | **/###/^ |

mean±SEM; Statistic: *,^ <0.05; **, ##,^^ <0.01; ***, ### <0.001; ****, #### <0.0001, ns: non-significant

Statistic: WT vs. *Pkd1* $^{RC/RC}$ (star, *); *Pkd1* $^{RC/RC}$ 3mo vs. 6mo or 3mo vs. 9mo (number sign, #); *Pkd1* $^{RC/RC}$ 6mo vs. 9mo (circumflex, ^)

#### Supplemental Table 2 | Analysis of the macrophage and epithelial cell population in mouse wildtype and ADPKD1 kidneys.

| | CD64$^+$ (%live) | | | APN$^+$/EpCAM$^+$ (%live) | | |
|---|---|---|---|---|---|---|
| C57Bl/6J | WT | *Pkd1* $^{RC/RC}$ | *Statistics* | WT | *Pkd1* $^{RC/RC}$ | *Statistics* |
| 3mo | 1.75±0.17 | 2.16±0.23 | ns | 85.93±1.73 | 89.12±0.79 | ns |
| 6mo | 2.93±0.44 | 4.54±0.36 | ns/#### | 77.43±2.47 | 74.99±2.04 | ns/#### |
| 9mo | 1.99±0.21 | 4.57±0.82 | */ns/ns | 84.93±1.55 | 67.30±2.55 | ****/####/^ |
| 129S6/SvEVTac | WT | *Pkd1* $^{RC/RC}$ | *Statistics* | WT | *Pkd1* $^{RC/RC}$ | *Statistics* |
| 3mo | 0.88±0.08 | 2.68±0.42 | *** | 89.45±0.73 | 83.97±1.50 | ** |
| 6mo | 1.23±0.19 | 3.70±0.45 | ***/ns | 88.34±1.20 | 85.89±1.06 | ns/ns |
| 9mo | 2.09±0.45 | 7.12±0.79 | **/###/^^ | 86.97±0.94 | 73.44±1.90 | **/##/^^ |
| BALB/cJ | WT | *Pkd1* $^{RC/RC}$ | *Statistics* | WT | *Pkd1* $^{RC/RC}$ | *Statistics* |
| 3mo | 0.55±0.10 | 3.86±0.53 | *** | 93.70±0.67 | 87.68±1.26 | ** |
| 6mo | 0.70±0.09 | 3.56±0.71 | ***/ns | 89.68±1.21 | 81.35±2.29 | */ns |
| 9mo | 1.28±0.41 | 3.10±0.49 | */ns/ns | 88.48±3.08 | 71.40±7.28 | ns/#/ns |

mean±SEM; Statistic: *,^ <0.05; **, ##,^^ <0.01; ***, ### <0.001; ****, #### <0.0001, ns: non-significant

Statistic: WT vs. *Pkd1* $^{RC/RC}$ (star, *); *Pkd1* $^{RC/RC}$ 3mo vs. 6mo or 3mo vs. 9mo (number sign, #); *Pkd1* $^{RC/RC}$ 6mo vs. 9mo (circumflex, ^)

**Supplemental Table 3 | PKD histomorphometric analyses of various ADPKD1 models with either genetic *Pd-l1* loss or anti-PD-1 treatment.**

| | Cyst Size ($\mu m^2$) | Cyst Number/$mm^2$ | Fibrotic Index (%) |
|---|---|---|---|
| C57Bl/6J *Pkd1* $^{RC/RC}$ | | | |
| *Cd274* $^{+/+}$ | 21,775±2,310 | 15.49±1.45 | 14.60±2.25 |
| *Cd274* $^{-/-}$ | 18,891±1,289 | 16.53±1.35 | 12.70±1.20 |
| BALB/cJ *Pkd1* $^{RC/RC}$ | | | |
| *Cd274* $^{+/+}$ | 30,496±5,502 | 12.77±1.086 | 14.32±1.02 |
| *Cd274* $^{-/-}$ | 29,404±5,926 | 13.32±0.75 | 11.75±2.11 |
| 129S6/SvEVTac *Pkd1* $^{RC/RC}$ | | | |
| IgG2a | 64,099±8,355 | 9.02±0.75 | 17.03±0.74 |
| α-PD-1 | 57,198±5,850 | 9.00±0.66 | 15.47±1.45 |
| C57Bl/6J *Pkd1* $^{RC/-}$ | | | |
| IgG2a | 136,407±16,515 | 6.27±0.50 | 26.64±3.52 |
| α-PD-1 | 161,353±23,387 | 6.10±1.24 | 26.52±2.87 |

mean±SEM; Mann-Whitney Test for all pairwise comparisons was non-significant.

**Supplemental Table 4 | Correlation data of adaptive immune cell phenotypes with PKD parameters in 3-month-old BALB/cJ *Pkd1*$^{RC/RC}$ mice part of the combination immune checkpoint blockade trial.**

| | %KW/BW | Cystic Index | Fibrotic Index |
|---|---|---|---|
| $CD8^+$, $CD44^+$/$CD69^+$ (%CD45) | | | |
| Pearson *r* | -0.387 | -0.402 | -0.436 |
| P value | 0.042 | 0.034 | 0.020 |
| $CD8^+$, $Ki67^+$ (%CD45) | | | |
| Pearson *r* | -0.373 | -0.314 | -0.342 |
| P value | 0.051 | 0.103 | 0.074 |
| $CD4^+$, $FoxP3^+$ (%CD45) | | | |
| Pearson *r* | 0.204 | 0.127 | 0.178 |
| P value | 0.299 | 0.521 | 0.365 |
| $CD8^+$, $CD44^+$/$CD69^+$ or $Ki67^+$ (%CD45) | | | |
| Pearson *r* | -0.400 | -0.390 | -0.424 |
| P value | 0.035 | 0.040 | 0.025 |
| $CD8^+$, $CD44^+$/$CD69^+$ or $Ki67^+$ (%CD45) (positive variable); CD4+, FoxP3+ (%CD45) (negative variable) | | | |
| Pearson *r* | -0.456 | -0.371 | -0.445 |
| P value | 0.015 | 0.052 | 0.018 |
